# Macroevolutionary dynamics in micro-organisms: generalists give rise to specialists across biomes in the ubiquitous bacterial phylum *Myxococcota*

**DOI:** 10.1101/2023.09.26.559479

**Authors:** Daniel Padfield, Suzanne Kay, Rutger Vos, Christopher Quince, Michiel Vos

## Abstract

Prokaryotes dominate the Tree of Life, but our understanding of the macroevolutionary processes generating this diversity is still limited. Habitat transitions are thought to be a key driver of prokaryote diversity, but we still know relatively little about how prokaryotes successfully transition and persist across environments, and how this varies between biomes and lineages. Here, we investigate biome transitions and specialisation in natural populations of a focal bacterial phylum, the *Myxococcota*, sampled across a range of replicated soils and freshwater and marine sediments in Cornwall (UK). By targeted deep sequencing of the protein-coding gene *rpoB*, we found >2000 unique *Myxococcota* lineages, with the majority (77%) being biome specialists and <5% able to live across the salt barrier. Discrete character evolution models revealed that biome specialists very rarely transitioned to specialising in another biome. Instead, generalists mediated transitions between biome specialists. Multistate hidden-state speciation and extinction models found variation in speciation rate across the tree, but this variation was independent of biome association and specialisation. Overall, our results help explain how microbes transition between biomes and are consistent with “the jack-of-all-trades” trade-off, where generalists suffer a cost in any individual environment, resulting in rapid evolution of niche specialists.

## Introduction

Understanding the ecological and evolutionary forces that structure prokaryote diversity across heterogeneous environments is a central objective in microbial ecology [1–3]. The extent to which different taxa are associated with different biomes, the frequency at which taxa transition between these biomes, and how this influences their diversification rate are not yet fully understood. One of the most drastic environmental transitions for both macro- and microorganisms is that between marine and terrestrial (land and freshwater) biomes, the so-called “salt barrier” [4]. Salinity is a major determinant in structuring microbial diversity, with distinct phylogenetic shifts observed over salinity gradients [5]. Transitions between marine and terrestrial biomes require substantial re-organisation of the proteome [6,7] and gains and losses of a multitude of genes and metabolic pathways [5,6,8–12]. Due to these adaptive challenges, microbe transitions across the salt barrier are thought to be rare [4,6].

Trade-offs between ecological specialisation strategies may explain the scarcity of successful transitions across the marine-terrestrial divide in prokaryotes. Generalist taxa that can live in both terrestrial and marine environments (and easily transition between them) are expected to be at a competitive disadvantage in any individual biome according to the classic “jack-of-all-trades, master of none” trade-off [13]. Apart from antagonistic pleiotropy, a generalist strategy could have other fitness costs, such as reduced evolvability [14]. In macro-organisms, evolutionary transitions between generalism and specialism are thought to occur in both directions, but are more commonly directed towards specialism [15,16]. Recent studies on prokaryotes that classified generalists or specialists based on their distribution across environments also found that evolutionary transitions are directed predominantly towards specialism, but additionally, that generalists possessed higher speciation rates [17,18]. These results highlight the key role generalists may play in colonising novel environments and generating microbial diversity [17,18].

However, these studies on prokaryotes did not apply some of the newly developed comparative phylogenetic methods that can better test whether variation in diversification rates is due to the focal trait (e.g. specialist or generalist) [19] and relied on the 16S rRNA gene marker, which, although representing the ‘gold standard’ in microbial ecology, offers relatively low genetic resolution and occurs in multiple (sometimes different) copies per genome [20]. The most commonly used alternative, metagenomic sequencing [6], likely misses rare taxa, which can affect the results of diversification analyses [21]. An alternative option is to use single-copy protein-coding genes, which are reliable proxies for whole-genome divergence [22] and with a high rate of evolution enables differentiation between even closely related taxa [23,24]. Moreover, amplicon sequencing targeting a subgroup of bacteria should allow exhaustive sampling and retrieval of taxa that would otherwise remain hidden.

Here, we use the *rpoB* gene to amplify Myxobacteria (previously classified as the δ-Proteobacterial Order *Myxococcales* but recently proposed to form the Phylum *Myxococcota* [25,26]), best known for their social development into multicellular fruiting bodies, large genomes, and prolific production of secondary metabolites [27,28]. Myxobacteria have long been known to live across a wide range of terrestrial habitats [29], but in the last two decades, they have also been shown to be ubiquitous components of marine [30] and freshwater [31] sediments. We sequenced the *rpoB* gene in samples collected from different marine and terrestrial environments replicated across Cornwall (U.K.) and, as has been done in a recent study [32], classified samples not based on a set of abiotic measurements (e.g. pH or salinity), but based on 16S rRNA community composition, reasoning that this reflects both the abiotic environment (i.e. ASVs in a sample share abiotic affinities) and biotic environment (i.e. ASVs co-occur in a sample because they interact with each other). The presence/absence of *Myxococcota* ASVs across biomes was used to classify each as a specialist or generalist, and we used modern comparative methods to investigate their macroevolution. Models of discrete character evolution revealed that generalists form ‘evolutionary bridges’ between biome specialists and act as sources of specialist lineages, with transitions predominantly directed towards specialists. Using the state-dependent speciation and extinction (SSE) framework, we found that diversification rates across the phylogeny varied, but were not associated with biomes or degree of specialisation. Our results demonstrate how generalists successfully transition between biomes, but then relatively rapidly evolve into specialists, likely due to the “jack-of-all-trades master-of-none” trade-off.

## Results

### Extensive regional phylogenetic diversity of the phylum Myxococcota structured across three main biomes

We used targeted sequencing of a ∼225 base pair (bp) region of the *rpoB* gene to uncover Myxobacterial diversity across the county of Cornwall (U.K.). Fifteen predefined, more or less distinct habitats were sampled across six broad locations with a view to maximising ecological and phylogenetic diversity, including freshwater, estuarine, and marine sediments, and soils associated with different vegetation or land uses (Figure 1A; Table S1). After prevalence filtering, 2,621 unique *Myxococcota* amplicon sequence variants (ASVs) were identified, compared to a total of only 153 *Myxococcota* ASVs retrieved from 16S sequencing (a 17-fold increase). We also saw increased diversity and relative abundance of *Myxococcota* in individual samples. In the *rpoB* dataset, we identified an average (mean) of 239 ASVs per sample (minimum = 1, maximum = 789) and the relative abundance of *Myxococcota* was 0.14 (minimum = <0.001, maximum = 0.47), compared to an average diversity of 42 (one sample had zero *Myxococcota*) and an average proportion of 0.02 in the 16S data, representing a seven-fold increase in *Myxococcota* sequences. Rarefaction curves demonstrated that diversity was sequenced to sufficient depth across all samples (Figure S1), and assigning taxonomy using the lowest common ancestor method (see Methods) resulted in 97% of all ASVs being assigned to at least family level. All seven named families in the *Myxococcota* were retrieved, alongside 16 unidentified families, demonstrating that our primers had phylum-wide coverage.

**Figure 1.**
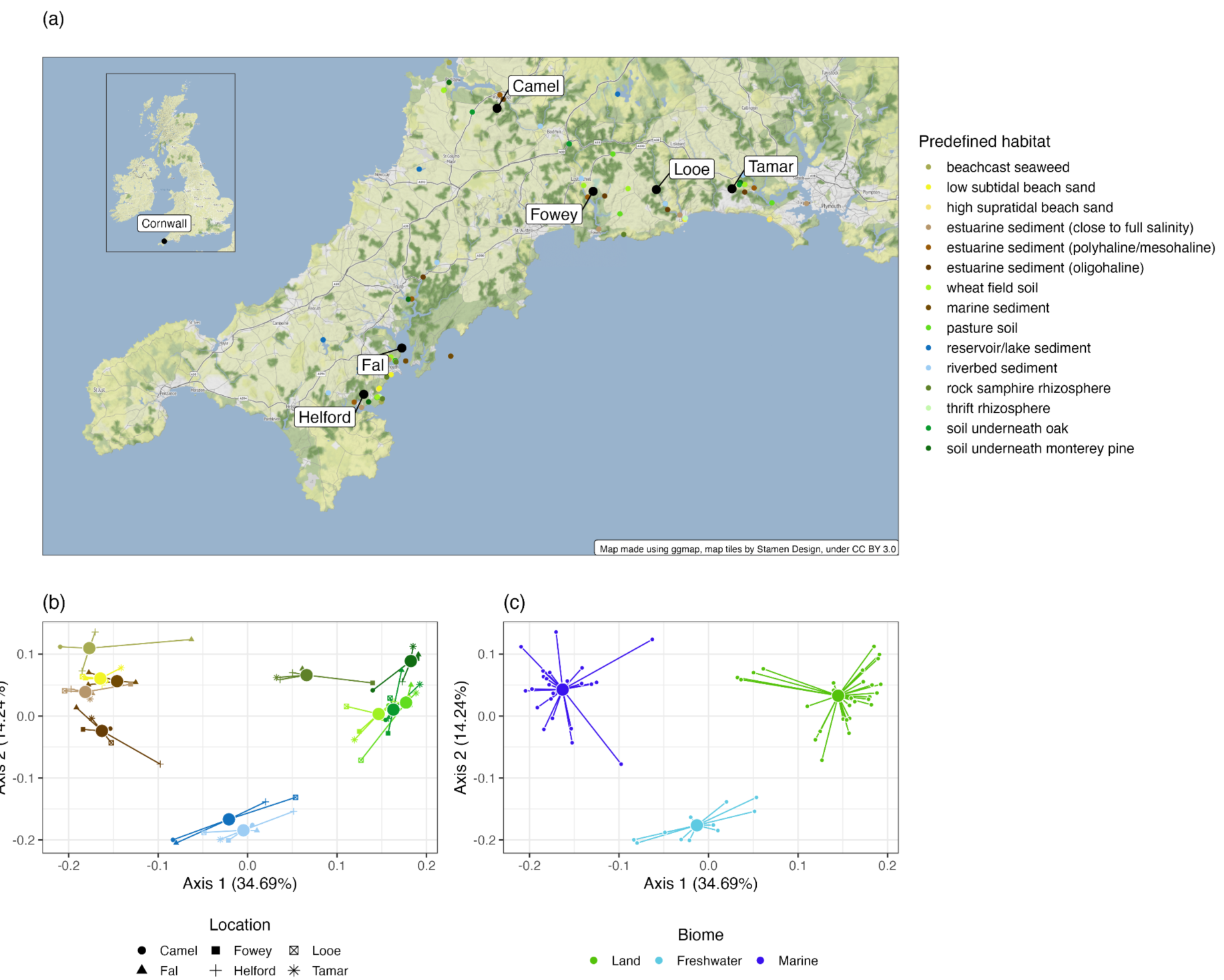
Predefined habitats and biome clusters from our sampling sites across Cornwall, UK. (a) Sampling locations of different predefined habitats across six locations in Cornwall in the south west of the UK. (b) Principal Coordinate (PCoA) plot of samples based on the weighted-Unifrac distance of the 16S data, with samples coloured by their habitat. Samples cluster together based on habitat (different colours), not location (different shapes). (c) PCoA plot of samples based on the weighted-Unifrac distance of the 16S data with samples coloured by their assignment into a habitat cluster based on medoid clustering. The best clustering resulted in three clusters: Land (green), Marine (dark blue), and Freshwater (light blue). In (a) large black points represent broad sampling sites, and small points represent specific sampling sites. In (b) and (c), each small point is an individual sample, large points are the positions of centroids of that group of samples, and lines connect individual samples to the group centroid.

To determine whether predefined habitats indeed differed, we looked for an overall effect of habitat and location on community composition as quantified by 16S rRNA sequencing (Figure 1b). Our predefined habitats explained a significant amount of variation in microbial community composition (PERMANOVA, F_14,53_=13.17, R^2^=0.76, p=0.001), whereas geographical location, as expected [33,34], did not (PERMANOVA, F_5,53_=1.14, R^2^=0.024, p = 0.278) (Figure 1b). To determine which predefined habitats differed significantly in community composition, we ran multiple pairwise permutational ANOVAs (see Methods) and removed three predefined habitats (high supratidal beach sand, thrift rhizosphere, and estuarine sediment (polyhaline/mesohaline)) that were not significantly different from the other predefined habitats. This left us with 12 predefined habitats, each with a significantly different community composition. As having 12 states for our observed trait for comparative analyses is computationally intractable, we used k-medoid clustering to calculate the optimal number of clusters based on the principal coordinate analysis of community composition that corresponded to the three main biomes: freshwater (11 samples), marine (25), and land (27)(Figure 1c).

To assign biome preference to each *Myxococcota rpoB* ASV, we compared their observed prevalence across all biomes to that expected by chance (accounting for the unequal numbers of samples in each biome). Most ASVs (77%) were associated with only one of the three major biomes and were designated as either freshwater (738), land (704) or marine (568) specialists. Generalism was designated when an ASV was present in multiple biomes at proportions equal to or exceeding those expected by chance and was rarer than specialism (23% of all ASVs). Only six ASVs were designated ‘true’ generalists present in all biomes, five ASVs were classified as land + marine generalists, 112 as freshwater + marine generalists, and 488 ASVs as freshwater + land generalists. Therefore, ignoring the latter generalist category, only 123 ASVs (<5%) occurred in both saline and non-saline environments, which is in line with our expectation that the salinity boundary is challenging to cross [4,6,35]. This principal finding of biome specialists being most common (and generalists capable of straddling the salt barrier being rare) was consistent across a range of OTU cut-offs (from ASVs to 91% similarity)(Figure S2).

We constructed an ultrametric phylogeny of all *Myxococcota* ASVs (Figure 2a), constraining the tree structure based on seven named family-level clades identified in a recent multi-gene phylogeny (Figure 2a inset) [26]. Of all ASVs, 73.6% were assigned to these seven families, with the remaining tips being unconstrained. We determined how biome associations were linked to the *Myxococcota* phylogeny in multiple ways. First, clustering of our *Myxococcota rpoB* sequencing based on the weighted Unifrac distance - which considers the phylogenetic proximity of species - demonstrated that freshwater, marine, and land samples had distinct *Myxococcota* composition (Figure S3). Second, we found strong phylogenetic signal of biome preference using two different methods: the D-statistic and Pagel’s lambda (Figure S4). The D-statistic is based on the sum of sister-clade differences in a given phylogeny, and is equal to one if the observed trait has a random distribution and to zero if the observed trait is as clumped as if it had evolved by Brownian motion, but values can fall outside of this range. In our phylogeny, D-statistic values [36] for the binary (yes or no) ability to live in freshwater, marine, or land biomes were 0.31, −0.20, and 0.04 respectively, with all of these being significantly different from a random distribution (p < 0.001) indicating significant phylogenetic clumping. Pagel’s lambda measures phylogenetic dependence of an observed trait, with a value of 1 indicating the trait has evolved by perfect Brownian motion, and lower values indicating that other factors, unrelated to phylogenetic history, have an impact on trait evolution. The estimate of Pagel’s lambda [37] for biome preference was 0.86, meaning that a large proportion of the variation in biome preference is explained by the phylogeny. The level of phylogenetic signal did not change systematically across different OTU cut-offs and was unaffected by tree size (after bootstrapping the tree at each OTU cut-off to the same size) (Figure S4). The clustering results and strong phylogenetic signal indicate biome preference is conserved across the *Myxococcota* phylogeny, meaning that closely related *Myxococcota* are likely to be ecologically similar.

**Figure 2.**
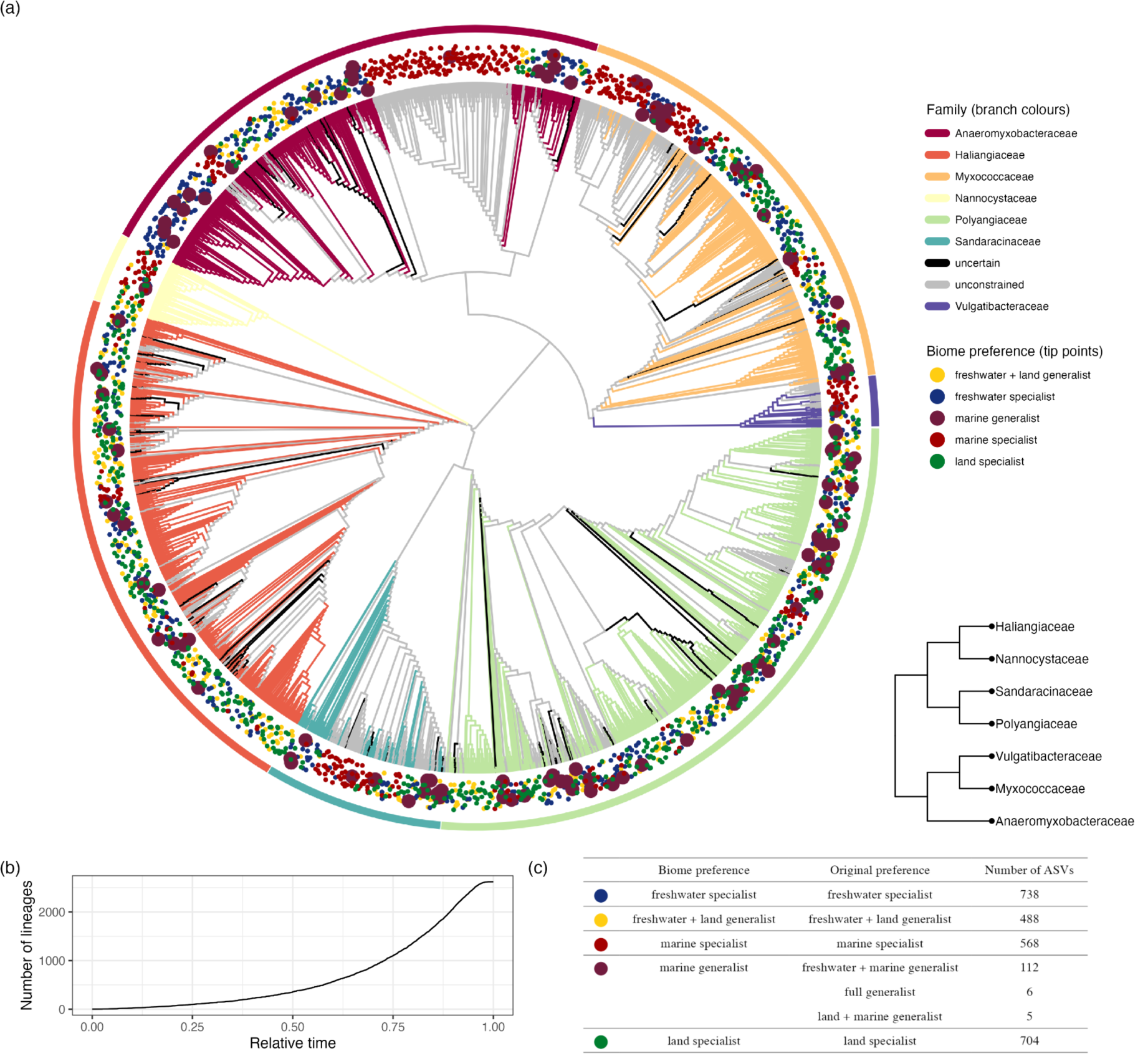
ASV-level constrained phylogeny of Myxococcota. (a) Ultrametric phylogenetic tree of Myxococcota from the rpoB sequencing. We constrained our phylogenetic tree using a recent Myxococcota multi-gene phylogeny (inset a) and allowed ASVs not assigned to one of the seven families to be unconstrained. Branch colours represent different family taxonomic assignments that we constrained when making the phylogeny; black represents ASVs without a family assignment, and grey represents unconstrained ASVs. Points around the tips of the tree represent biome preference of each ASV. Large points allow easier visualisation of marine generalits as they are the least common. (b) Lineage through time plot for the accumulation of new ASVs through relative time. (c) Table showing the classification and abundance of different biome preferences of *Myxococcota*.

### Biome transitions are mediated by generalists

We next tested whether differences in transition rates between specialists and generalists drove the uneven distribution of ASVs across biome specialists and generalists. As it is difficult to fit comparative phylogenetic models when distributions across states are extremely uneven and when some states have low numbers, we collapsed the three biome preferences with the smallest numbers of ASVs (marine + land generalist, freshwater + marine generalist, and full generalist) into a single preference of “marine generalist” (Figure 2c). We tested ten Markov models to study discrete character evolution and explore the transitions between biome preferences in the *Myxococcota* through evolutionary time. We fitted four hypothesis-driven models that restricted some transitional pathways: all-rates-different (ARD), symmetric (SYM), equal rates (ER), and stepwise (SW). The ARD model assumes all transitions are possible and all rates can differ. The SYM model assumes all transitions are possible, but rates to and from any pair of biome preferences are equal, and the ER model assumes all transitions are possible but all occur at the same rate. The SW model assumes that an intermediate generalist state is needed to move to a new specialisation (i.e. evolution from marine specialist to land specialist requires a marine + land generalist step first), but all allowed rates can differ. The remaining six models were simplifications of the ARD model where we iteratively set transitions with the lowest rates to zero (see Methods). A custom ARD model of just 11 transitions (of a possible 20) was best supported (AIC weight = 0.39, Figure 3a, Table S2), while the estimated transition rates were qualitatively similar among the four best-supported models that cumulatively had an AIC weight of 0.87 (Figure 3).

**Figure 3.**
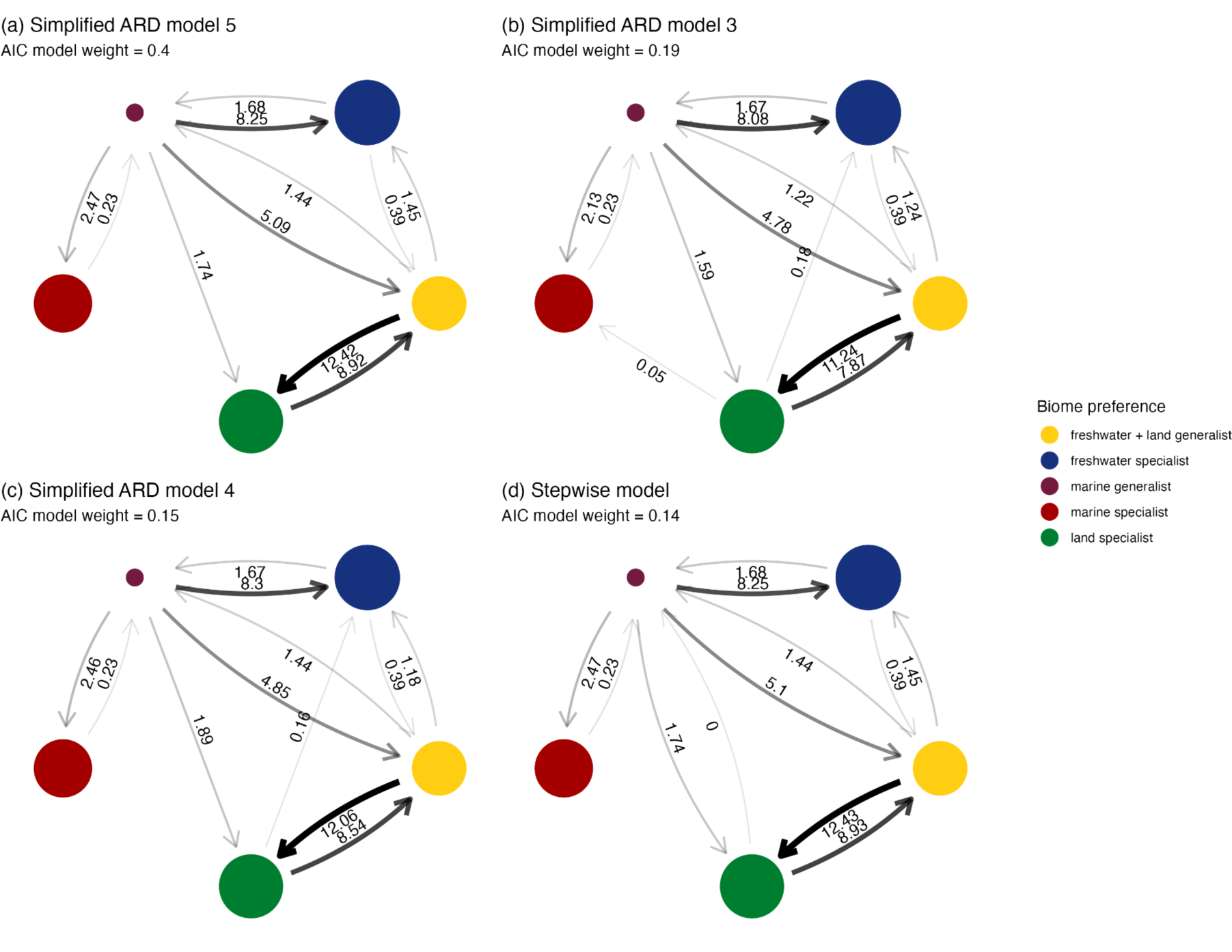
Transition rates between biome preferences for the four best-supported models of discrete character evolution. The three best-supported models (a-c) were simplifications of the ARD (all rates different model) where low transition rates were removed. The fourth best-supported model (d) was the stepwise model, which did not allow direct transitions between specialist states or marine specialists and freshwater + land generalists. The radius of circles is proportional to the number of ASVs in each biome preference. The size of the arrows is proportional to the transition rate. Transition rates are labelled to two decimal places.

The parameter estimates of the transition models revealed several key patterns in the evolution of biome preference in the *Myxococcota*. First, it is very rare for specialists of one biome to shift directly to specialising in another biome. The best-supported model does not support any specialist-to-specialist transitions (out of potentially six transitions; Figure 3a), and across the four best-supported models, only the transition from land specialist to freshwater specialist is estimated to occur (Figure 3b,c), and at a very low rate. Instead, marine generalists and freshwater + land generalists were the best connected (with seven and six transitions, respectively, compared to a maximum of four for a biome specialist), acting like stepping stones through which biome specialists evolve (Figure 3). Second, generalists are less stable than specialists, with transition rates away from the more generalist state exceeding those towards these states (Table 1). For instance, transition rates away from marine generalists are more than five times higher than all rates towards it combined, and freshwater + land generalists have the second highest ratio of rates directed towards compared to away from. In contrast, all specialist states are stable, with transition rates into freshwater, marine, and land specialists being 79%, 91%, and 37% higher than those away from these states (Table 1). Moreover, when looking at individual pairs of transitions, transition rates away from the more generalist state were always higher than rates towards it. This pattern is consistent across all four best supported models (Figure 3). Third, marine specialists are extremely stable and the most evolutionarily isolated of all biome preferences, with transition rates both towards and away from this state being the lowest compared to all other biome preferences (Table 1). Fourth, transitions between land specialists and freshwater + land generalists are widespread, indicating that species can easily transition between these biome preferences. We can exclude the possibility that freshwater + land generalists simply represent land specialists transiently present in freshwater sediments due to runoff, as the best supported Markov model indicates that freshwater + land generalists are more connected to other states than to land specialists. Our bootstrapping approaches, where we (a) subsampled 80% of the tree or (b) subsampled the tree to have the same number of tips within each biome preference and then re-fitted the best supported Markov model, gave qualitatively equivalent results (Figures S5 and S6). Specifically, transitions away from biome generalists were higher than transitions away from biome specialists; marine generalists were the least stable and marine specialists were the most stable biome preference.

**Table 1.**
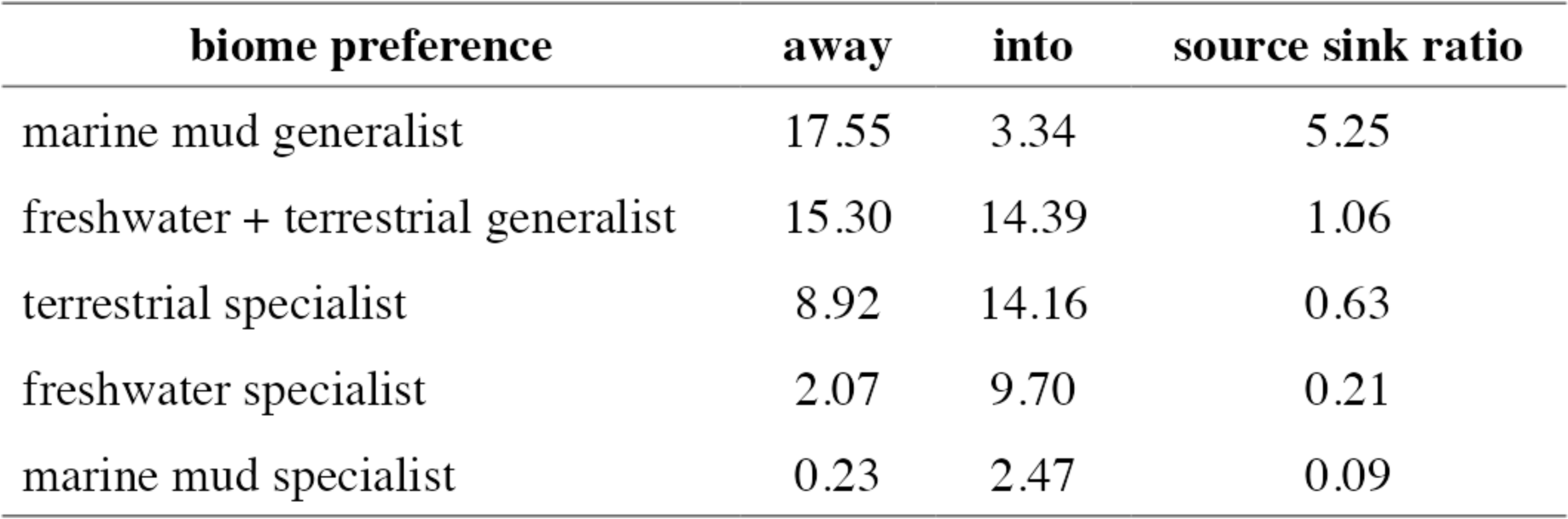
Total transition rates to and from each biome preference.

| **biome preference** | **away** | **into** | **source sink ratio** |
| --- | --- | --- | --- |
| marine mud generalist | 17.55 | 3.34 | 5.25 |
| freshwater + terrestrial generalist | 15.30 | 14.39 | 1.06 |
| terrestrial specialist | 8.92 | 14.16 | 0.63 |
| freshwater specialist | 2.07 | 9.70 | 0.21 |
| marine mud specialist | 0.23 | 2.47 | 0.09 |

### Heterogeneity in the diversification and speciation of Myxococcota does not vary between biome specialists and generalists

We used Bayesian Analysis of Macroevolutionary Mixtures (BAMM) [38] to detect shifts in diversification rates across our *Myxococcota* phylogeny (Figure 4a). Speciation rates across the phylogeny generally decreased over time, while extinction rates remained relatively stable, resulting in a net decrease in diversification rate (Figure 4b). More important than the average rates across the whole tree, we detected heterogeneity in the diversification rate across the tree, with an average of 12 shifts in the diversification regime (95% credible intervals: 9-18)(Figure 4c). Supporting this, a model with 12 rate shifts in the diversification regime had the highest posterior probability (0.19) and was strongly supported over models with fewer or more regime shifts (closest Bayes Factor difference = 34). However, the evidence for any one of the 993 detected shift configurations was very weak, with the highest percent probability for a single shift configuration being only 0.19%. Consequently, we calculated the best overall shift configuration and estimated the diversification rate through time averaged over the whole tree and for subsets of the tree at the nodes where core shifts occurred during this configuration (see Methods and Figure 4a). Immediately after a core shift in diversification rate, rates spiked before receding back towards the global average (Figure 4a). In summary, we find evidence of heterogeneity in speciation rates across the *Myxococcota*, but it is impossible to ascertain precisely where, how often, and with what magnitude these shifts are occurring in the phylogeny. To test whether heterogeneity in diversification rate was associated with biome preference, we fitted a set of multi-state dependent speciation and extinction models (MuSSE) that allow diversification rates to vary with biome preference while accounting for the transition rates between these states [39]. A MuSSE model with only state-dependent speciation rates was selected over models with (a) both state-dependent speciation and extinction, (b) state-dependent extinction only, and (c) no state-dependent speciation or extinction (AIC weight = 0.98). This result is consistent with the BAMM analysis that found that heterogeneity in diversification rates was mainly driven by speciation rate. The MuSSE model showed that marine generalists had a significantly higher speciation rate than the other biome preferences. However, SSE analyses must be interpreted with caution as they rest on the assumption that rate heterogeneity is associated with variation in the measured trait state (e.g. biome preference) [19,40].

**Figure 4.**
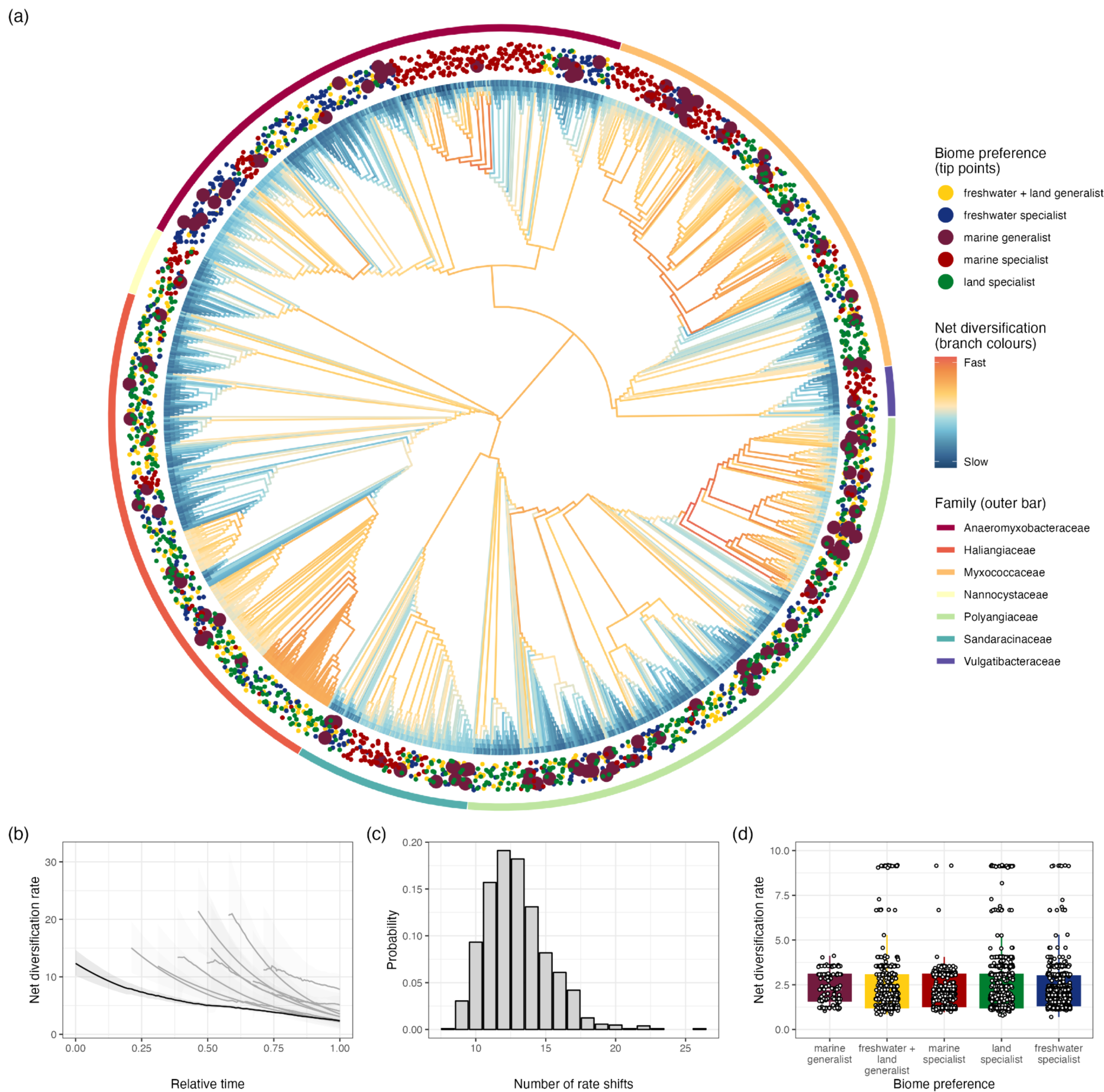
Rate heterogeneity in the diversification of *Myxococcota*. (a) Ultrametric phylogenetic tree of *Myxococcota* with rates of net diversification inferred using BAMM. Branch colours represent the diversification rate (warmer colours = higher rates). Points around the tips of the tree represent biome preference of each ASV, and bars around the tree represent the family of each clade. (b) Rate-through-time plot showing how net diversification decreases over evolutionary time. The black line represents the average across the whole tree; the grey lines represent the rate-through-time on parts of the tree where core rate shifts were identified. Shaded regions represent 95% confidence intervals. (b) Posterior distribution of the number of rate shifts inferred using BAMM. (d) Variation in tip-specific diversification rates inferred using BAMM across biome generalists and specialists.

To address this shortcoming, we fitted several models containing “hidden” (or concealed) traits, so-called Hidden State-dependent Speciation and Extinction Models (HiSSE) [19,41,42]. To reduce the number of estimable parameters, we fixed the transition rates to those estimated from the best Markov model. Doing so had little impact on the diversification rate estimates (Pearson’s correlation coefficient between speciation rates with- and without fixing some transition rates = 0.99), and the correlation between transition rates estimated from the MuSSE model and the Markov model was strong (Pearson’s correlation coefficient = 0.95). We refitted the MuSSE model with only state-dependent speciation and compared it to several null models. First, concealed-trait-dependent (CTD) models [19] with two, three, or four concealed states, in which rates of speciation were allowed to vary across lineages modulated by hidden states but not by biome preference. Second, a MuHiSSE model [42] allowing diversification rate variation owing to both hidden variation and biome preference. The CTD4 model (a concealed-trait-dependent model with four hidden states) performed best by far (AIC weight ∼ 1) and had diversification rates broadly in line with the values found in the original MuSSE, with one hidden state having a high speciation rate (15.54, compared to 12.7 for marine generalists), two intermediate rates (2.59 and 5.72, compared to 3.88 for marine specialists), and one low rate (0.57, compared to 1.96 for freshwater specialists). Across different estimated sampled fractions of diversity, the concealed-trait dependent models were favoured over MuSSE or MuHiSSE models, with CTD4 performing best at an assumed sample fraction of 0.75 and CTD3 performing best at a sample fraction of 0.5 (Table S3). In summary, after accounting for hidden states, there is no evidence for differences in diversification rates between biome specialists or generalists. In line with this result, a phylogenetic regression on the tip-specific evolutionary rates estimated from BAMM found no significant difference in diversification rates between biome preferences (all contrasts between biome preferences had p > 0.05, Figure 4d).

## Discussion

In this study, we used an ecologically and geographically explicit, replicated sampling design to explore biome transitions and specialisation in the macroevolutionary history of the *Myxococcota*. Specifically, we used 16S sequencing to cluster predefined habitats into three clusters corresponding to the main freshwater, land, and marine biomes (Figure 1c). Most *Myxococcota* ASVs were biome specialists, with less than 5% of ASVs able to live across the salt barrier (Figure 2c). We used models of discrete character evolution to demonstrate how specialisation evolves and biome transitions occur. Generalists mediate transitions between biomes and then rapidly evolve into specialists, which, while not evolutionary dead ends, generally have much lower transition rates away from them than generalists (Figure 3, Table 1). Finally, we performed analyses investigating variation in diversification rates across the *Myxococcota* and found shifts in diversification rate (Figure 4), but these shifts were not attributable to biome preference or specific *Myxocococcota* clades (Table 2). Our results revealed that the much rarer biome generalists mediate transitions between biomes, with transition rates substantially higher away from, rather than into, generalist states, consistent with previous findings based on 16S rRNA data [17,18,43]. There was very little statistical support for transitions between biome specialists, suggesting that biome generalists successfully transition between biomes, after which they rapidly evolve to specialise on one specific biome. Crucially, this result was replicated across all biomes, with rates into freshwater/land/marine specialists always being higher than rates in the opposite direction towards a generalist (Figure 3a). In this way, generalists act like ‘stepping stone’ lineages through which microbes transition between biomes before evolving specialisation, somewhat similar to work that found brackish water biomes act like ‘stepping stone’ environments mediating marine-terrestrial transitions [6]. Not all biome specialists evolve in the same way, with marine specialists being the most evolutionarily isolated, with transition rates into and away from this biome specialisation the lowest of any state (Table 1). This might reflect more constrained pathways of adaptation from saline to non-saline environments, but also more constrained dispersal routes: migration into the marine environment might be more frequent than the other way around thereby offering more potential for colonising taxa to adapt to this environment.

**Table 2.** Model comparison of multi-state and concealed trait diversification rate models.

| Model | Number of estimated parameters | Log Likelihood | AIC | AIC weight |
| --- | --- | --- | --- | --- |
| CTD4 | 17 | -1,289.80 | 2,613.60 | 1 |
| CTD3 | 10 | -1,313.83 | 2,647.65 | 0 |
| MuHiSSE | 13 | -1,339.34 | 2,704.68 | 0 |
| CTD2 | 5 | -1,376.03 | 2,762.07 | 0 |
| MuSSE | 6 | -1,719.65 | 3,451.29 | 0 |

Our analyses did not uncover differences in diversification rates between generalists and specialists associated with different biomes. Historically, specialisation has been considered an evolutionary dead end, which may result in lower speciation rates and higher extinction rates. Still, recent analyses have demonstrated that diversification rate is often independent of specialisation in macro-organisms. In microbes, studies have shown mixed results, with two studies finding that generalists had (much) higher diversification and speciation rates than specialists [17,18], while the other demonstrated the opposite pattern [43]. However, none of these studies used Hidden State Speciation and Extinction models, which can account for unknown (hidden) traits that may affect diversification rate [19,40], meaning there is a high risk of false positives in these analyses. In line with this, our MuSSE analysis found that marine generalists had higher speciation rates than other biome preferences, but after using HiSSE models, the best-supported model was one where the diversification rate was independent of biome preference.

Similar to previous work investigating bursts in diversification rate in prokaryotes [44,45], we found evidence of rate shifts in diversification (speciation) rates across the phylogeny, but it was not possible to assign such bursts to specific taxa, biomes, or generalist/specialist strategies. Bursts in diversification rate are often interpreted as adaptive radiations, which occur when a single ancestral type encounters broad ecological opportunity, enabling diversification into a multitude of specialised types [46]. Although their capacity for dispersal means that adaptive radiations in prokaryotes are unlikely to arise via colonisation events of novel ecosystems, it may be that the uptake of novel traits through horizontal gene transfer (HGT) allows the colonisation of new niche space, which subsequently can be partitioned into different specialists. Going forward, combining targeted amplicon sequencing data with (metagenome-assisted) whole-genome data [2,6] is needed to characterise the role of HGT in evolutionary transitions and its mechanistic impact on ecological specialisation [2].

In summary, we present the first work - to our knowledge - that looks at the macroevolution of both biome transitions and specialisation in prokaryotes simultaneously and is amongst the first to apply SSE methods to an observed trait (biome preference) with more than two states. In doing so, we demonstrate how generalists allow for successful transitions between biomes, including across the salt barrier, before rapidly giving rise to specialists. The exponential rise in microbial sequencing data will undoubtedly result in comparative phylogenetics being increasingly used to understand prokaryote evolution, but using these methods for this purpose remains challenging [47]. First, the lack of a universally accepted (operational) species definition in prokaryotes impedes the estimation of global diversity and inferring phylogenetic species trees. Second, while targeted short amplicon sequencing allows for better sequencing of the microbial diversity in a sample, it can be hard to estimate robust phylogenetic trees, whereas the opposite is true for metagenomic data. These issues can affect the results of diversification rate analyses [21,47], but as the majority of prokaryotic diversity is currently unculturable, there is no other way to study the macroevolution of bacteria. Our approach here was to cross-validate our analyses as thoroughly as possible, compare the relative fit of multiple models to understand how biome preference evolved, and compare relative differences in parameter estimates rather than concentrating on absolute values. Increased collaboration between comparative phylogeneticists and microbial ecologists is needed to uncover macroevolutionary dynamics in prokaryotes.

## Methods

### Environmental sampling

We sampled 15 predefined habitats in August 2020 replicated across six drowned river valleys (‘rias’) in Cornwall (United Kingdom): Helford, Fal, Fowey, Looe, Tamar and Camel (Figure 1a). Predefined habitats were riverbed sediment, reservoir/lake sediment, pasture soil, wheat field soil, soil underneath oak, soil underneath monterey pine, rock samphire rhizosphere, marine sediment, low subtidal beach sand, high supratidal beach sand, beachcast seaweed, estuarine sediment (close to full salinity), estuarine sediment (polyhaline/mesohaline), estuarine sediment (oligohaline), and thrift rhizosphere. The replication levels differed among habitats due to practical limitations, resulting in 73 samples in total (Table S1). Each of the soil or sediment samples consisted of multiple subsamples taken from an area of approximately 0.25m^2^ to minimise stochastic variation (as it is likely that individual subsamples will contain a variety of (micro)niches [48]). Soil samples were taken as shallow as possible after removing leaf litter and were sieved to remove debris. Each sample was stored in two 50mL falcon tubes and frozen upon return to the lab at −70°C.

### DNA extraction and 16S sequencing

DNA extractions were carried out according to the Qiagen DNeasy PowerSoil kit handbook (1104560 HB-2257-001). A 10-minute incubation at 70°C after the lysis step was included to increase DNA yield. DNA quantity was verified using a picogreen assay (qubit HS DNA kit)(Invitrogen), purity was assessed using nanodrop 260:280 ratios, and integrity was evaluated using a 1% agarose gel. A 251bp conserved fragment in the V4 hypervariable region of the 16s rRNA gene was targeted using N515f and N806r primers [49] with a pool of indexed primers suitable for multiplex sequencing with Illumina technology. Sequencing was performed using an Illumina MiSeq 500-cycle V2 Kit by the University of Exeter Sequencing Service. After the first sequencing run, four samples had very low depth and were resequenced. Sequencing adapters and any bases below a phred score of Q22 were removed, alongside any reads <150bp, using ‘*Cutadapt*’ (v4.4) [50]. Reads were processed in R (v4.2.2) using the packages *‘dada2*’ [51] and ‘*phyloseq’* [52]. As error rates differ between sequencing runs, we estimated trimmed reads, estimated error rates, and inferred and merged sequences separately. While processing the first sequencing run, we trimmed the first 10bp off and truncated all reads at 225bp for both the forward and reverse samples. For the four re-sequenced samples, we trimmed the first 10bp off the forward and reverse reads and then truncated forward reads at 265bp and reverse reads at 225bp. We then merged the two sequence tables (which joined together any amplicon sequence variants (ASVs) present across sequencing runs), removed chimeric sequences, and assigned taxonomies to ASVs using the SILVA database (v138.1) [53]. We estimated a phylogeny using ‘*fasttree*’ [54]. Any ASVs that (a) were over 250bp in length, (b) had not been assigned to at least Phylum level, (c) appeared in <5% of all samples and (d) had a total abundance of <200 across the whole dataset were removed. Overall, this left 6030 individual ASVs across 73 samples encompassing the 15 habitats that were included in downstream analyses, with an average of 58,681 reads per sample, a minimum of 12,570 reads and a maximum of 174,902 reads.

### rpoB amplicon primer design and sequencing

Group-specific primers targeting the Myxobacteria (GTDB Phylum *Myxococcota*) were designed using the R package DECIPHER [55]. Firstly, all genomes assigned to the phylum *Myxococcota* from the NCBI [56] and GTDB (r202) [25] databases were downloaded using ‘*ncbi-genome-download*’ [57] to extract the *rpoB* gene sequence. We removed identical sequences, kept only sequences between 3900bp and 4400bp in length, and ensured there was only a single copy of *rpoB* per genome (keeping the sequence closest to the median length of the gene). Finally, we removed five sequences that aligned especially poorly to the others. The remaining 158 sequences were aligned using ‘*DECIPHER::AlignTranslation()*’, resulting in a 4641bp alignment. Outgroup sequences were chosen by re-rooting the GTDB phylogeny (r202: https://data.gtdb.ecogenomic.org/releases/release202/202.0/bac120_r202.tree) to the origin of the *Myxococcota* and selecting the 3000 accessions that had the shortest distance to this node (i.e. the bacteria most closely related to the *Myxococcota*). The genomes for these accessions were downloaded, and the *rpoB* gene sequence aligned as described above, but in addition, we removed sequences that had a median distance (from the other outgroup sequences) of over 0.4 and a distance from a reference *Myxococcus xanthus* DK 1622 sequence of over 0.35. This resulted in 164 non-*Myxococcota* sequences and a 4689bp alignment.

Both alignments were combined using ‘*AlignProfiles()*’ to create a 322 sequence, 5060bp alignment of *Myxococcota* and non-*Myxococcota* sequences. Primers were designed using ‘*DECIPHER::DesignPrimers()*’. No selective primers for long amplicons could be designed, so we limited our search to a predicted product size between 200 and 400bp. Several candidate primers were tested on genomic DNA of *Nannocystis exedens*, *Bradymonas sediminis*, and *Corallococcus coralloides* (purified gDNAs purchased from the Leibniz Institute DSMZ Braunschweig, Germany). We also tested these primers on gDNA extracted from a random sample of river sediment using our chosen purification method. The primer pair producing a single strong product for all test samples was selected from the candidate list. Our final primers for targeted *Myxococcota rpoB* sequencing were GCGATCAAGGAGCGCATG-F and CAGATGCGGCCGTAGTG-R. This primer set had a predicted amplicon size of ∼260bp, and was predicted to amplify 78% of the *Myxococcota* sequences in our alignment and only 5% of the non-*Myxococcota* sequences. We created phased primer pairs to sequence 63 samples; samples from high supratidal beach sand, thrift rhizosphere, and estuarine sediment (polyhaline/mesohaline) were removed as they did not differ in composition from the majority of other predefined habitats. Sequencing was done on an Illumina Novaseq on 28/09/2021 with paired-end 250bp reads by the Exeter Sequencing service. Primers were removed, and reads were dephased before being processed using ‘*dada2*’ and ‘*phyloseq*’.

First, forward and reverse reads were truncated at 200bp. The Novaseq sequencing run returned binned quality scores, which meant the estimated error rates at the highest quality score were higher than those at intermediate quality scores. To overcome this, we enforced monotonicity to the error model by changing the arguments of the loess model to have a span equal to 2 and weights equal to the log-transformed total counts of nucleotides (https://github.com/benjjneb/dada2/issues/1307). We then inferred and merged sequences, constructed a sequence table, and assigned taxonomy using a reference database of all *rpoB* sequences in the GTDB database (r202). This pipeline resulted in a 222bp *rpoB* amplicon, 494,114 unique ASVs and a mean read number of 1,570,304 (minimum=220,231, maximum=7,427,083). We filtered this dataset to solely retain ASVs assigned to any of the *Myxococcota* phyla in the r202 GTDB database (*Myxococcota*, *Myxococcota_A*, and *Myxococcota_B*).

To cross-validate the naive Bayesian classifier implemented in *dada2*, the taxonomy of all sequences identified as *Myxococcota* was also assigned using lowest common ancestor (LCA) algorithms as implemented by ‘*MMSeqs2’* [58]. After building a custom database from the GTDB *rpoB* fasta file, taxonomy was assigned using the default LCA algorithm (*mmseqs taxonomy --lca-mode 3*), selecting the most specific taxonomic label that had at least 95% support (*--majority 0.95*) of the −log(E-value) weights (*--vote-mode 1*). Additional arguments set were: assigning taxonomy to nucleotide sequences (*--search-type 3*), returning all lineage information in the output (*--tax-lineage 1*), and disabling pre-filtering query ORFs (*--orf-filter 0*). This resulted in the removal of 76 ASVs (0.17%) not assigned to *Myxococcota*. Prevalence filtering removed all *rpoB* ASVs occurring in fewer than four samples and with a total abundance of fewer than 100 reads. After these filtering steps, there were 2,621 individual ASVs, and samples had a mean read number of 87,948 (minimum=28, maximum=340,355).

To test whether there was a level of phylogenetic relatedness of *Myxococcota* that best explained variation in biome preference, we clustered our dataset at 1% increments of OTU similarity from ASV level (100%) to 91%, with an additional cut-off at 97.7% as that was previously identified as a suitable species boundary cut-off for *rpoB* [23]. The 2,621 unique sequences were aligned using ‘*DECIPHER::AlignSeqs()*’ using a guide tree, and the distance matrix was calculated using ‘*DECIPHER::DistanceMatrix()*’. For each OTU similarity cut-off, we clustered the sequences from the distance matrix using ‘*DECIPHER::TreeLine()*’ and used ‘*speedyseq::merge_taxa_vec()*’ [59] to merge clusters into single OTUs (using the name and sequence for the most abundant ASV in the cluster to represent the new clustered OTU).

For each clustered dataset, we estimated a phylogenetic tree using *raxml-ng* (v1.1.0) [60]. We used a recent multi-gene phylogenetic tree of the *Myxococcota* [26] to create a constraint tree, ensuring that any ASVs assigned to the families *Myxococcaceae*, *Vulgatibacteraceae*, *Anaeromyxobacteraceae*, *Polyangiaceae*, *Sandaracinaceae*, *Nannocystaceae,* and *Haliangiaceae* were placed within the same clade, and relationships between families were fixed based on the topology of the multi-gene tree. ASVs not assigned to one of these families were left unconstrained. We used the GTR + gamma model and ran 20 tree searches (10 random and 10 parsimony-based starting trees), and the best tree was chosen based on the best-scoring topology. For the best tree of each clustered dataset (100-91% OTU similarity), the tree was rooted manually in FigTree [61] by finding the split between the two Classes (*Myxococcia* or *Polyangia*) specified in the constraint tree. Trees were made ultrametric in R using ‘*ape::chronopl()*’ [62] with a smoothing parameter of 10.

### Statistical Analyses

#### Analysing microbial community composition and clustering samples into biomes

To test whether our predefined habitats differed in community composition, we employed both supervised and unsupervised clustering analyses on the relative abundances of the 16S ASVs using weighted Unifrac distance [63]. First, we ran a permutational ANOVA to test whether habitat or location had significant impacts on community composition using ‘*vegan::adonis2()*’ [64] with 9999 permutations. Following this, we ran pairwise permutational ANOVAs between all pairs of predefined habitats to test which were significantly different from each other. This was done by subsetting the data into pairwise combinations of habitats, running a permutational ANOVA on each subset, and extracting the *R^2^* value and *p*-value, which was adjusted using the false discovery rate (fdr) method [65]. The only non-significant contrasts involved samples from high supratidal beach sand, thrift rhizosphere, and estuarine sediment (polyhaline/mesohaline), which were removed from subsequent analyses.

Predefined habitats were then clustered into broad biomes using unsupervised learning, with the dissimilarity matrix created from multidimensional scaling of the weighted-Unifrac distance matrix used as the input and limiting the maximum dimensions of the space of the matrix to only include positive eigenvalues. K-medoid and hierarchical clustering methods were used to estimate the number of clusters that best grouped the data using two approaches. First, we used k-medoid clustering using ‘*cluster::clusGap()*’ [66] at every level of possible clustering (from 1 to 12 - the number of predefined habitats). The optimal number of clusters was calculated using the gap statistic and their standard deviations, using Tibshirani’s recommendation [66,67]. Second, we used k-means clustering and ‘*NbClust::NbClust()*’ [68], which calculates 30 indices and recommends the optimal number of clusters using the majority rule. We also used the gap statistic and the majority rule approaches to determine the optimal number of clusters using hierarchical clustering, where we used the ward method in ‘*clusGap()*’.

All four combinations of clustering (k-medoid and hierarchical) and methods to determine optimal cluster numbers (gap statistic and majority rule) assigned samples to three clusters (freshwater, land, and marine). The single difference was that both hierarchical clustering methods assigned one sample of beach seaweed to the land cluster, whereas all beach seaweed samples were assigned to the marine cluster using k-medoid clustering methods. As it makes sense for all samples within a predefined habitat to be clustered within the same biome, the samples were assigned to clusters using the k-medoid clustering method, with the gap statistic and majority rule approaches giving identical results.

#### Assigning biome preference to Myxococcota ASVs

The presence/absence of each *Myxococcota* ASV across the three biomes (freshwater, marine or land) was used to assign biome preference. Any ASV that was only present in a single biome was designated as a biome specialist. For any ASV present in two or three biomes, we employed a bootstrapping approach to assign biome preference. Specifically, we created a bootstrapped presence dataset for each ASV by sampling their observed presence across samples 100 times with replacement to calculate their proportional presence across biomes. This process was repeated 1000 times for each ASV to create a distribution of observed biome preference proportions. We then compared these observed proportions to those based on the number of samples in each biome (land = 0.44, marine = 0.38, freshwater = 0.18), akin to habitat availability. For every ASV, if 97.5% of the observed use estimates in any given biome were below the expected proportion given its availability, we assumed it did not have an affinity for that biome. Consequently, ASV biome preference consists of all the biomes where the ASV was present at a level at least as high as expected by each biome’s availability. This approach meant that seven different biome preferences were possible (freshwater specialist, marine specialist, land specialist, freshwater + marine generalist, marine + land generalist, freshwater + land generalist and full generalist (i.e. land, freshwater and marine). Biome preference was assigned separately to each ASV at each OTU similarity cut-off.

#### Estimating phylogenetic signal in biome preference

Phylogenetic signal of biome preference for the *Myxococcota* was estimated in two ways. First, we calculated the D-statistic, which estimates the amount of phylogenetic signal of binary traits [36]. As biome preference has seven potential values, we constructed three new traits based on an ASV’s ability to live in any of the three biomes (freshwater: yes or no; land: yes or no; marine: yes or no). We then calculated the D-statistic separately for each trait using ‘*caper::phylo.d()*’ [69]. Second, we estimated Pagel’s lambda by using ‘*geiger::fitDiscrete()*’ [70] for the biome preference trait, using an equal rates model. We estimated phylogenetic signal at each OTU similarity cut-off; as the size of the tree can impact estimates of phylogenetic signal, we recalculated D and lambda at each OTU similarity cut-off after randomly subsetting the tree to 350 tips (as the size of the smallest tree was 353 tips) and repeated this 100 times to visualise the variation around this estimate. There was no systematic variation in phylogenetic signal across the OTU similarity cut-off range (Figure S4), so the original ASV dataset was chosen for all downstream analyses. There was also minimal impact of tree size on the estimates of phylogenetic signal (Figure S4).

#### Investigating the evolution of biome preference using models of discrete character evolution

We modelled the evolution of biome preference using Markov models. We used ‘*diversitree::fit_mk()*’ [39] which can handle multi-state traits and estimate transition rates among different states. As the numbers of ASVs that were full generalists or marine + land generalists were extremely low, we merged these with freshwater + marine generalists to create a marine generalist group. We fitted four hypothesis-driven models that restricted some transitional pathways: all-rates-different (ARD), symmetric (SYM), equal rates (ER), and stepwise (SW). As our trait is simply an association, these are not the only biologically plausible models. Consequently, we also performed iterative model simplification on the ARD model to set the lowest transition rate to zero until AIC stopped decreasing. For the first simplified model, we set four transitions that were less than 0.001 to 0 and then set the single smallest transition rate to zero for each subsequent model simplification. We then compared all models using Akaike Information Criterion (AIC) weights [71]. For the best-supported model, we calculated a “source-sink ratio” by dividing the sum of the transition rates into a biome preference by the sum of the transition rates away from the same biome preference. We estimated the uncertainty in transition rates using two bootstrap approaches. First, we subsampled the tree to 80% of its full size and re-fitted the best-supported model. Second, we subsampled each group to have the same number of ASVs (123) and re-fitted the best model. We did both approaches for 1000 iterations and then calculated mean estimates and 95% confidence intervals for transition rates and “source-sink ratios”.

#### Exploring heterogeneity in diversification rates of Myxococcota using BAMM

We used BAMM (Bayesian Analysis of Macroevolutionary Mixtures) to estimate speciation and extinction rates and identify rate shifts in net diversification across our *Myxococcota* phylogeny [38]. BAMM uses reversible-jump Markov chain Monte Carlo sampling to explore shifts in macroevolutionary regimes, assuming they occur across branches of a phylogeny under a compound Poisson process. It explicitly explores diversification rate variation through time and among lineages. Priors for BAMM were generated using the R package ‘*BAMMtools*’ [38] and the expected number of transitions was set to 500 to aid convergence [72]. We ran four MCMC chains for 30,000,000 generations, allowing chain swaps every 1000 generations and saving output every 20,000 generations. We assessed convergence by calculating the effective sample size (ESS) of the log-likelihood and the number of shifts of the results after a burn-in period of 30% (ESS values >200 are indicative of good convergence). We also checked that the posterior of the number of transitions differed from the prior by using ‘*BAMMtools::plotPrior()*’.

The best overall model (number of rate shifts) from our BAMM analysis was chosen by selecting the model with the highest Bayes factor relative to the null model, which has zero rate shifts. We calculated the credible shift set - the ranked set of distinct shift configurations that accounts for 95% of the posterior probability of the data - for our BAMM analysis. This returns the number of core shifts, defined as those that contribute appreciably to our ability to model the data. In contrast, non-core shifts are simply shifts we would expect to sometimes happen under the prior distribution for rate shifts across the tree. In our case, all shift configurations had very low probability (the best having a posterior probability of 0.0019). This is expected in some datasets with large numbers of taxa as there are simply too many parameters in the model to allow a single shift configuration to dominate the credible set. Consequently, we extracted the shift configuration using ‘*maximumShiftCredibility*’ that maximises the marginal probability of rate shifts along individual branches, similar to the maximum clade credibility tree in phylogenetic analysis. We used phylogenetic generalised least squares regression using ‘*nlme::lme()*’ [73] to determine whether tip-specific net diversification rates estimated from BAMM were associated with biome preference. Although disagreements exist about what these estimates represent [74], this analysis helped complement the state-dependent diversification analyses. In this model, tip-specific net diversification rate was the response variable, biome preference was the predictor, and the phylogenetic tree was used to create a correlation matrix to control for the effect of phylogeny. Post-hoc comparisons between individual pairs of biome preferences were done to see if any biome preferences had significantly different tip-specific net diversification rates.

#### State-dependent diversification analysis and parameterisation

We used multi-state-dependent speciation and extinction (MuSSE) models [39] to determine whether rate heterogeneity is associated with biome preference. In these models, a lineage’s speciation or extinction rate depends on biome preference, and transitions between biome preferences were limited to those from the best supported transition matrix from the Markov models. We first used ‘*diversitree::fitmk()*’ to compare models where (1) both speciation and extinction were associated with biome preference, (2) only speciation was associated with biome preference, (3) only extinction was associated with biome preference, and (4) neither speciation nor extinction was associated with biome preference (constant-rates model). Models were compared using AIC weights.

It is possible that the SSE model could be supported over a constant-rates model just because it allows for variation in speciation (or extinction) rate across the tree [19,40]. Consequently, we fitted models where diversification rates depend on an unknown (hidden or concealed) trait using the R package ‘*secsse*’ [19]. For all models, we estimated a single extinction rate and fixed transitions between biome preferences to those estimated from the best Markov model to limit the number of estimable parameters. We fitted four different models: (1) a MuSSE model with no hidden states, (2) a MuHiSSE model that allowed for both state-dependent and hidden state speciation rates, (3) a concealed trait diversification model (CTD) with two hidden states (CTD2) (4) a CTD model with three (CTD3) states, four (CTD4), or five (CTD5) hidden states. For models including hidden states, transitions to and from the same hidden state were allowed to differ (i.e. 1A -> 1B ≄ 1B -> 1A), and dual transitions were disallowed (i.e. could not move hidden and measured traits at once). Models were compared using AIC scores and AIC weights. Statistical support for biome preference affecting diversification rates is found when the AIC score of the model in which speciation (or extinction) differs across biome preferences is higher than that in which rates depend on an unknown (CTD model) and a constant-rate model.

We used several different initial parameter sets to circumvent local optima during likelihood optimisation with ‘*secsse*’. The first set of parameters were the estimates of speciation and extinction from a birth-death model fit to the branching times and transition rates from the best-supported Markov model using ‘*DDD::bd_ml()*’ [75]. For transitions between hidden states, the initial start value was the mean of the transition rates from the best supported Markov model. We then created a grid of all combinations of starting values for half and double these initial values (27 different combinations) and calculated the log likelihood of the model using these estimates given the data using ‘*secsse::secsse_loglik()*’. We then chose starting parameters with the six highest initial log-likelihood values to fit to the data using ‘*secsse::secsse_ml()*’, choosing the highest likelihood of the six starting points to compare across models. We re-ran the model-fitting process at three different levels of sampling fraction (1, 0.75, and 0.5) to determine how sensitive our conclusions are to the assumption that we have sampled all the *Myxococcota* diversity in the samples. All models were fitted with a log-likelihood penalty of 0.05 to prevent unrealistically high parameter estimates and to aid in model fitting.

#### Overview of open source software used

All R scripts used elements of the suite of packages known as the ‘*tidyverse*’ [76], all phylogenetic trees were plotted using *‘ggtree’* [77], all figures were made using ‘*ggplot2*’ [78], and all tables were created using ‘*flextable*’ [79]. Specific R packages are referenced in their relevant section, and scripts for most parts of the analysis and to recreate all the plots created in the manuscript are available.

## Acknowledgements

All authors acknowledge funding support from the National Environment Research Council (NERC; grant NE/T008083/1). C.Q. was supported by BBSRC Core Strategic Programme Grant (BB/CSP1720/1, BBS/E/T/000PR9818, and BBS/E/T/000PR9817). D.P. was also funded by a NERC Independent Research Fellowship (NE/W008890/1) We thank Sarah Ashton and Jane Phelps for administrative support, Paul O’Neill and everyone at Exeter Sequencing Service, and Rampal Etienne for providing advice on using secsse. This work would not have been possible without tools generously made available by other researchers.

## Data Availability

All raw sequencing data will be deposited on the European Nucleotide Archive. The code to recreate all analyses is publicly available on GitHub (https://github.com/padpadpadpad/myxo_diversification) and will be archived on Zenodo upon publication. The analysis code starts with the processed phyloseq objects created after the dada2 workflow.

---

**Table S1.** Summary of samples, sites, predefined habitats, and sequencing.

| sample number | site | location | latitude | longitude | predefined habitat | 16s sequencing | rpoB sequencing |
| --- | --- | --- | --- | --- | --- | --- | --- |
| 60 | Camel | Doom Bar by Hawker's Cove | 50.56247 | -4.944736 | beachcast seaweed | yes | yes |
| 28 | Fal | Swanpool Beach | 50.14070 | -5.076021 | beachcast seaweed | yes | yes |
| 70 | Fal | Flushing | 50.16126 | -5.056420 | beachcast seaweed | yes | yes |
| 72 | Helford | beach near Rosemullion Head | 50.10760 | -5.087167 | beachcast seaweed | yes | yes |
| 30 | Fal | Penryn river Flushing | 50.16445 | -5.071440 | estuarine sediment (close to full salinity) | yes | yes |
| 38 | Fowey | Bodinnick ferry | 50.33983 | -4.630327 | estuarine sediment (close to full salinity) | yes | yes |
| 16 | Helford | Helford Passage Beach | 50.09993 | -5.128802 | estuarine sediment (close to full salinity) | yes | yes |
| 23 | Looe | Millpool pond | 50.35870 | -4.459996 | estuarine sediment (close to full salinity) | yes | yes |
| 52 | Tamar | The Old Rowing Club Torpoint | 50.37396 | -4.193850 | estuarine sediment (close to full salinity) | yes | yes |
| 64 | Camel | Wadebridge, Guinaport Road | 50.51337 | -4.831025 | estuarine sediment (oligohaline) | yes | yes |
| 55 | Fal | Tresillian River, Audi Tresillian | 50.27536 | -4.999346 | estuarine sediment (oligohaline) | yes | yes |
| 37 | Fowey | Lerryn | 50.38409 | -4.617869 | estuarine sediment (oligohaline) | yes | yes |
| 14 | Helford | Gweek Bridge | 50.09638 | -5.208347 | estuarine sediment (oligohaline) | yes | yes |
| 20 | Looe | Watgate camping West Looe | 50.36583 | -4.485670 | estuarine sediment (oligohaline) | yes | yes |
| 53 | Tamar | field near Polbathic Village Hall | 50.38930 | -4.323641 | estuarine sediment (oligohaline) | yes | yes |
| 63 | Camel | Wadebridge, Bridge Bike hire | 50.51930 | -4.838130 | estuarine sediment (polyhaline/mesohaline) | yes | no |
| 10 | Fal | Malpas Road pontoon | 50.24595 | -5.023255 | estuarine sediment (polyhaline/mesohaline) | yes | no |
| 41 | Fowey | Saint Winnow | 50.38258 | -4.653494 | estuarine sediment (polyhaline/mesohaline) | yes | no |
| 15 | Helford | Port Navas | 50.10716 | -5.144270 | estuarine sediment (polyhaline/mesohaline) | yes | no |
| 47 | Tamar | end of Old Quay Lane | 50.39482 | -4.303952 | estuarine sediment (polyhaline/mesohaline) | yes | no |
| 19 | Helford | Maenporth | 50.12460 | -5.093505 | high supratidal beach sand | yes | no |
| 44 | Tamar | Tregantle Beach/Whitsand Bay | 50.35302 | -4.270939 | high supratidal beach sand | yes | no |
| 59 | Camel | Doom Bar by Hawker's Cove | 50.56247 | -4.944736 | low subtidal beach sand | yes | yes |
| 27 | Fal | Gylly Beach | 50.14412 | -5.066652 | low subtidal beach sand | yes | yes |
| 18 | Helford | Maenporth | 50.12570 | -5.091337 | low subtidal beach sand | yes | yes |
| 24 | Looe | East Looe Beach | 50.35282 | -4.449455 | low subtidal beach sand | yes | yes |
| 42 | Tamar | Tregantle Beach/Whitsand Bay | 50.35249 | -4.271751 | low subtidal beach sand | yes | yes |
| 65b | Fal | Outer Bizzies (West) | 50.16904 | -4.941416 | marine sediment | yes | yes |
| 66 | Fal | Outer Bizzies (East) | 50.16904 | -4.941416 | marine sediment | yes | yes |
| 67 | Fal | East Narrows | 50.16247 | -5.036010 | marine sediment | yes | yes |
| 68 | Fal | East Narrows | 50.16247 | -5.036010 | marine sediment | yes | yes |
| 57 | Camel | road to Tregella Farm | 50.52576 | -4.955687 | pasture soil | yes | yes |
| 2 | Fal | Trefusis Head | 50.16199 | -5.057825 | pasture soil | yes | yes |
| 40 | Fowey | near Valleybrook | 50.36016 | -4.585863 | pasture soil | yes | yes |
| 5 | Helford | Maenporth Road | 50.11345 | -5.096796 | pasture soil | yes | yes |
| 34 | Looe | Wayton Barn | 50.44024 | -4.600625 | pasture soil | yes | yes |
| 46 | Tamar | Wacken Quay | 50.37494 | -4.266530 | pasture soil | yes | yes |
| 65 | Camel | Porth Reservoir | 50.42000 | -5.007679 | reservoir/lake sediment | yes | yes |
| 69 | Fal | College Reservoir | 50.15516 | -5.129523 | reservoir/lake sediment | yes | yes |
| 9 | Fal | Argal Reservoir | 50.15207 | -5.133878 | reservoir/lake sediment | yes | yes |
| 12 | Helford | Stithians Reservoir | 50.19099 | -5.209554 | reservoir/lake sediment | yes | yes |
| 33 | Looe | Colliford Lake | 50.52035 | -4.590757 | reservoir/lake sediment | yes | yes |
| 56 | Camel | Camel River Dunmere Bridge | 50.47749 | -4.753194 | riverbed sediment | yes | yes |
| 54 | Fal | Tresillian River | 50.29501 | -4.970236 | riverbed sediment | yes | yes |
| 35 | Fowey | Tudor Bridge | 50.40752 | -4.666920 | riverbed sediment | yes | yes |
| 13 | Helford | Helford Stream | 50.11960 | -5.198810 | riverbed sediment | yes | yes |
| 22 | Looe | West Looe River | 50.37369 | -4.490573 | riverbed sediment | yes | yes |
| 51 | Tamar | River Tiddy | 50.41212 | -4.327172 | riverbed sediment | yes | yes |
| 71 | Fal | Flushing | 50.16126 | -5.056420 | rock samphire rhizosphere | yes | yes |
| 29 | Fal | Swanpool Beach | 50.14104 | -5.075885 | rock samphire rhizosphere | yes | yes |
| 39 | Fowey | Lansallos Cove | 50.33249 | -4.577888 | rock samphire rhizosphere | yes | yes |
| 73 | Helford | Rosemullion Head | 50.11167 | -5.085058 | rock samphire rhizosphere | yes | yes |
| 43 | Tamar | Tregantle Beach/Whitsand Bay | 50.35302 | -4.270939 | rock samphire rhizosphere | yes | yes |
| 62 | Camel | Padstow School sports field | 50.53603 | -4.945018 | soil underneath monterey pine | yes | yes |
| 11 | Fal | St. Clement/Malpas | 50.24548 | -5.030602 | soil underneath monterey pine | yes | yes |
| 31 | Fowey | Lanhydrock | 50.45368 | -4.692297 | soil underneath monterey pine | yes | yes |
| 17 | Helford | Bosveal car park | 50.10749 | -5.114093 | soil underneath monterey pine | yes | yes |
| 49 | Tamar | Saltash | 50.40236 | -4.333044 | soil underneath monterey pine | yes | yes |
| 61 | Camel | Atlantic Highway lane | 50.49642 | -4.896110 | soil underneath oak | yes | yes |
| 1 | Fal | Kilnquay Wood | 50.16315 | -5.058057 | soil underneath oak | yes | yes |
| 32 | Fowey | Lanhydrock | 50.45405 | -4.692600 | soil underneath oak | yes | yes |
| 6 | Helford | Maenporth Road | 50.11400 | -5.093901 | soil underneath oak | yes | yes |
| 21 | Looe | West Looe river bank | 50.37369 | -4.490573 | soil underneath oak | yes | yes |
| 50 | Tamar | Saltash, Bag Lane | 50.39924 | -4.335000 | soil underneath oak | yes | yes |
| 3 | Fal | Flushing Beach | 50.16126 | -5.056420 | thrift rhizosphere | yes | no |
| 7 | Helford | Nansidwell | 50.11475 | -5.091429 | thrift rhizosphere | yes | no |
| 25 | Looe | East Looe Beach (rocks on the east side) | 50.35444 | -4.448103 | thrift rhizosphere | yes | no |
| 48 | Camel | Saint Germans | 50.39948 | -4.323441 | wheat field soil | yes | yes |
| 4 | Fal | Flushing | 50.16777 | -5.066961 | wheat field soil | yes | yes |
| 36 | Fowey | near Moorfield | 50.39811 | -4.662053 | wheat field soil | yes | yes |
| 8 | Helford | Maenporth Road | 50.11407 | -5.095674 | wheat field soil | yes | yes |
| 26 | Looe | Lanreath | 50.39391 | -4.568772 | wheat field soil | yes | yes |
| 58 | Tamar | road to Tregella Farm | 50.52539 | -4.956342 | wheat field soil | yes | yes |

**Figure S1.**
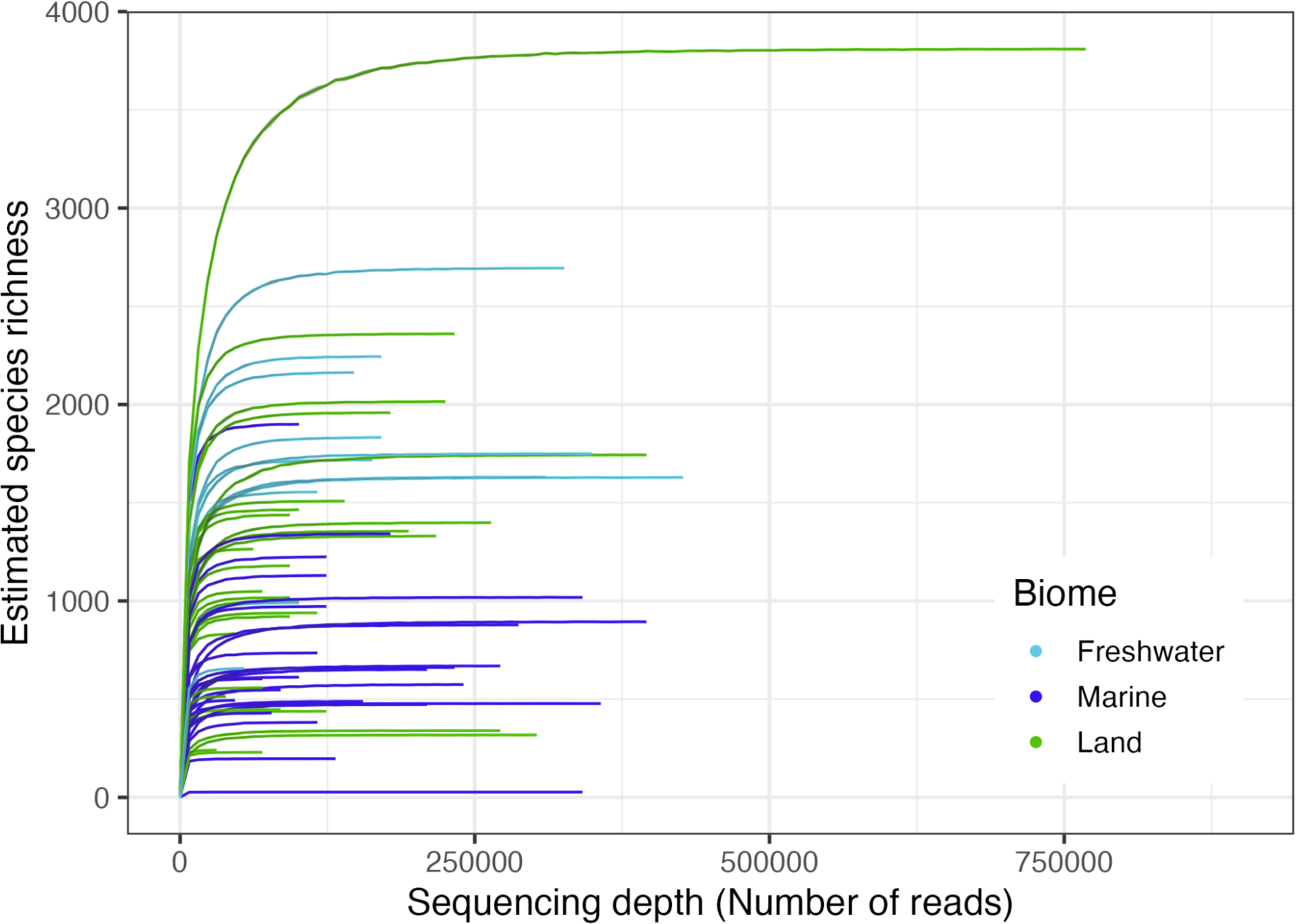
Rarefaction curves of rpoB sequencing. Rarefaction curves to check sufficient sequencing depth were estimated for each rpoB sequencing sample after filtering to keep only ASVs assigned to the *Myxococcota*. For each sample, rarefaction was constrained to the number of reads in each sample. The relationship between sequencing depth and estimated species richness has plateaued in all samples, indicating we have captured all the *Myxococcota* diversity in the sample (given the primers).

**Figure S2.**
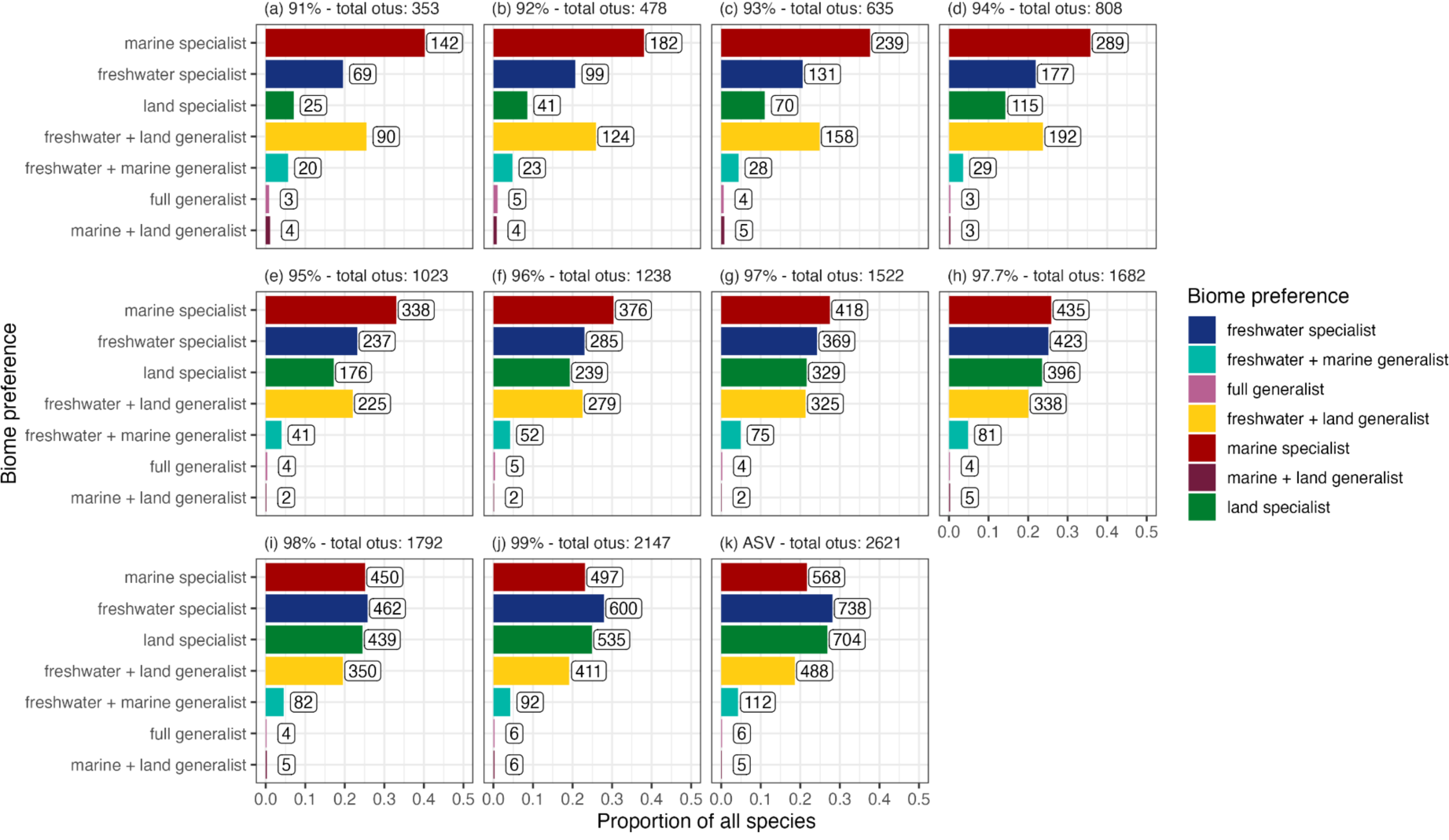
Total diversity and numbers of ASVs in each biome preference at different OTU similarity cut-offs. To look at whether phylogenetic signal in biome preference was highest at a specific OTU cut-off, we clustered the data from ASV (100%) to 91% OTU similarity, then performed prevalence filtering and assigned biome preference to each sequence variant. Proportions of sequence variants in each biome preference stay stable across OTU cut-offs: there are very few complete or marine + land generalists, and the dataset is mostly biome specialists.

**Figure S3.**
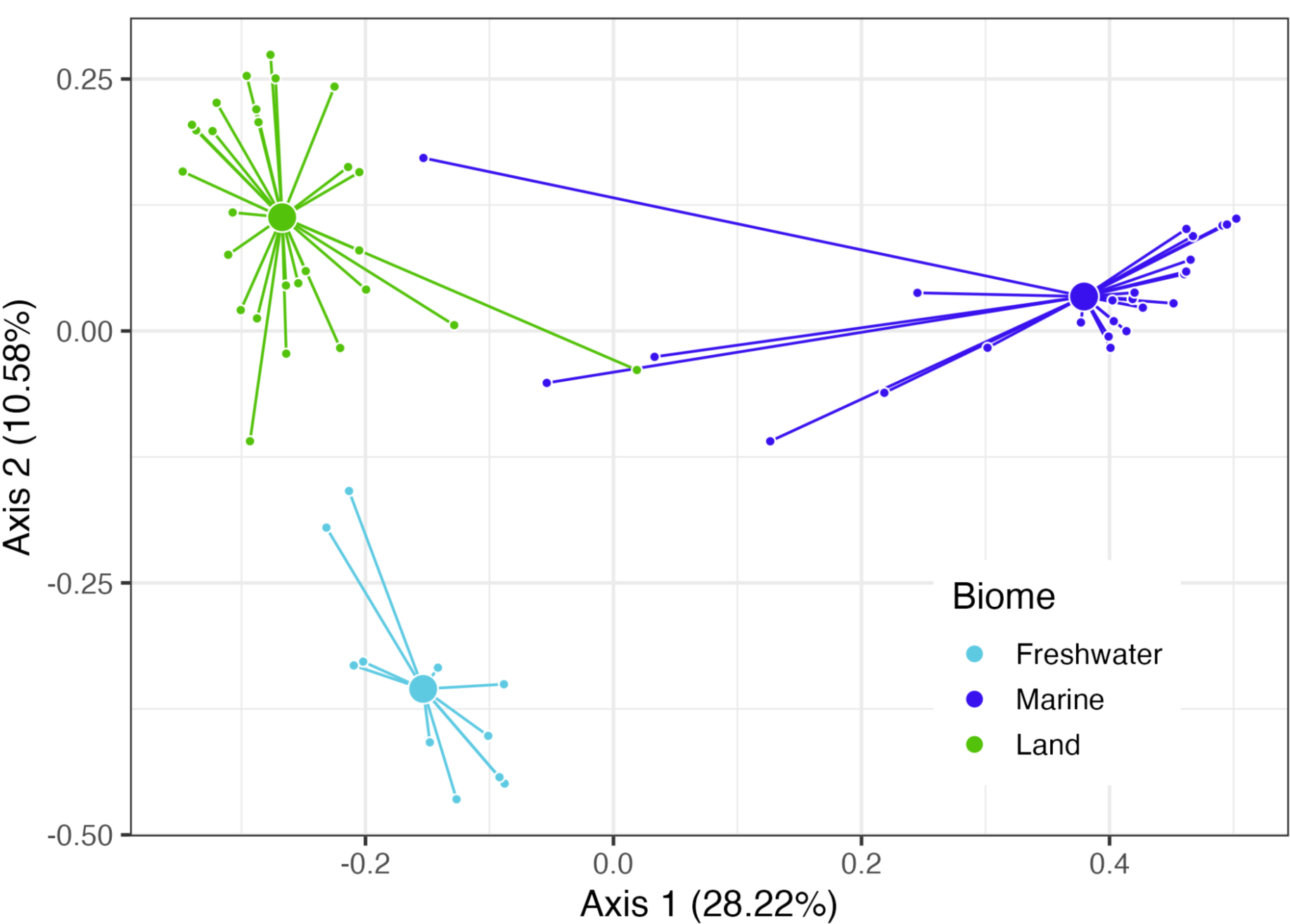
Principal Coordinate (PCoA) plot of samples based on weighted-Unifrac distance of the Myxococcota communities based on rpoB sequencing. There is significant clustering of *Myxococcota* communities, indicating that there the community composition is different in the different biomes. Each small point is an individual sample, large points are the positions of centroids of that group of samples, and lines connect individual samples to the group centroids.

**Figure S4.**
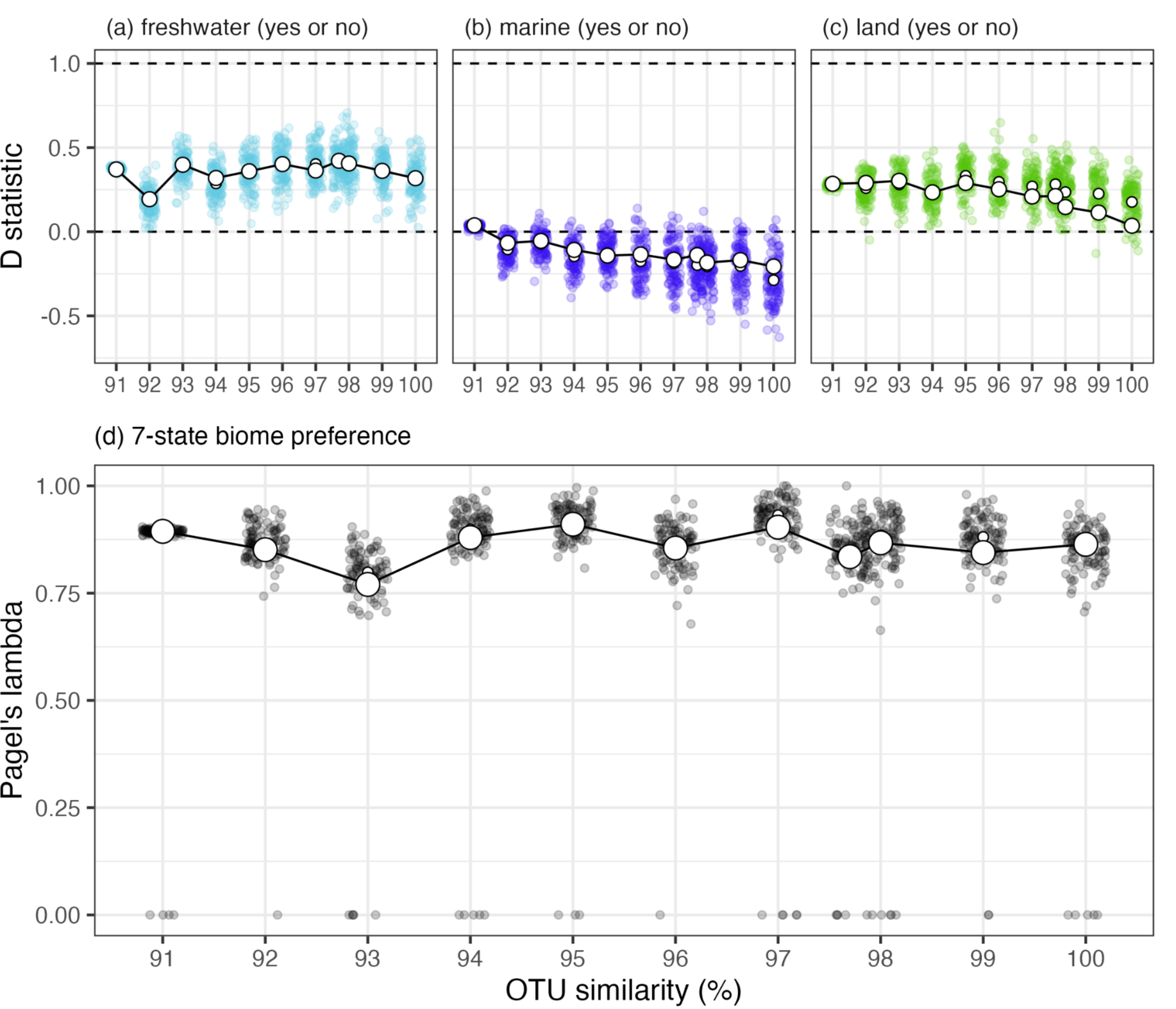
Phylogenetic signal of biome preference at different OTU similarity cut-offs. (a-c) D-statistic of binary traits of being able to live (or not) in each of the three broad biomes. (d) Pagel’s lambda estimate of the seven-state biome preference before collapsing marine + land, freshwater + land, and full generalists into a single preference. There was strong phylogenetic signal in all traits measured. In all panels, small black or coloured points represent individual bootstrapped estimates of the measure of phylogenetic signal where we have subsampled the tree to have the same size across OTU cut-offs (see Methods), small white points represent the average bootstrapped estimate. Large points represent the point estimate from the full tree at each cut-off.

**Figure S5.**
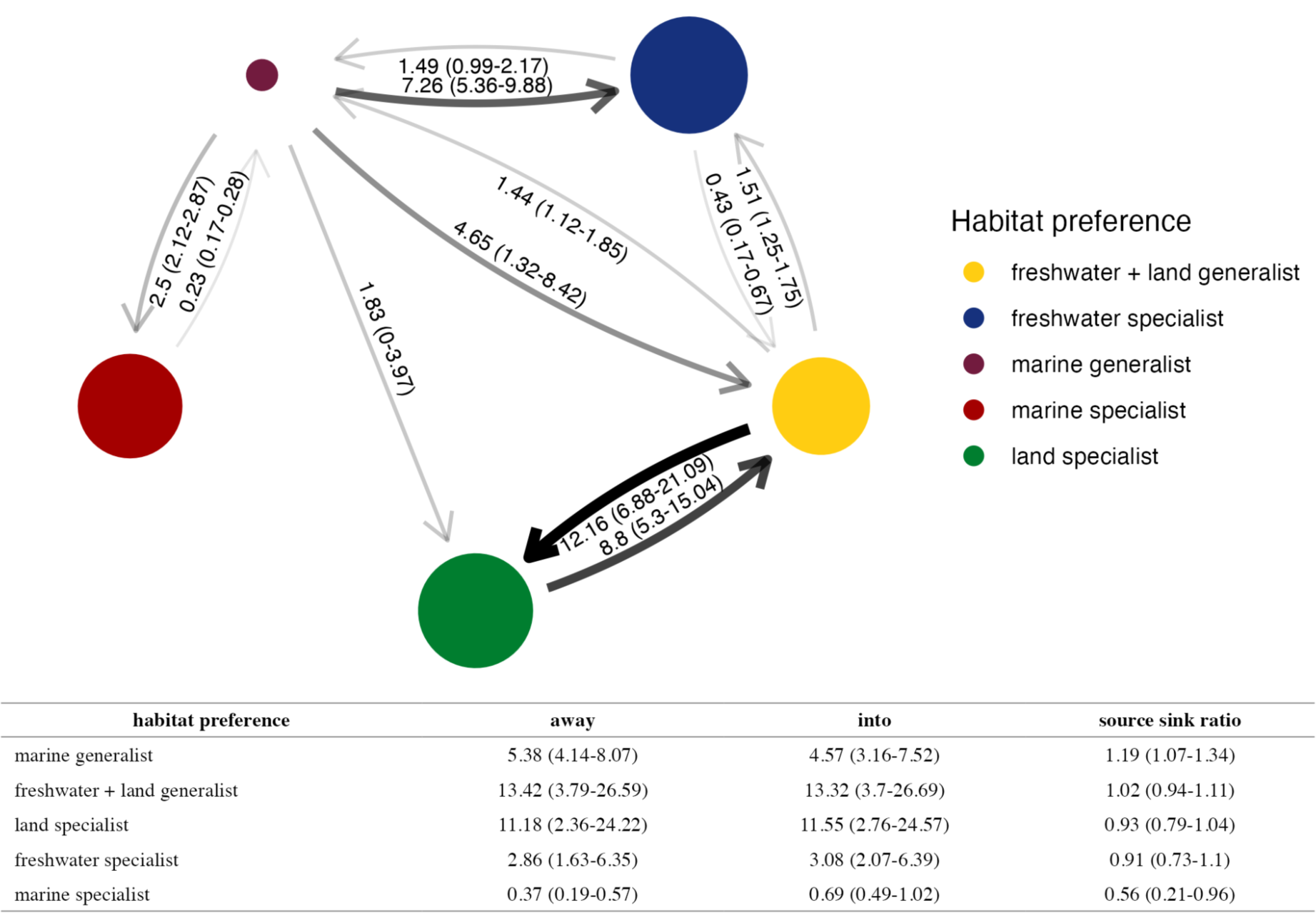
Bootstrapped (using random sampling) transition rates between biome preferences for the best-supported model of discrete character evolution. The best-supported model was a simplification of the ARD where low transition rates were removed. The table shows the total rates into and away from each biome preference. Bootstrapping was done by randomly sampling 80% of the tips of the tree and re-fitting the best-supported model. Mean estimates are presented, and values in brackets represent 95% confidence intervals. The radius of circles is proportional to the number of ASVs in each biome preference. The size of the arrows is proportional to the transition rate. All values are labelled to two decimal places.

**Figure S6.**
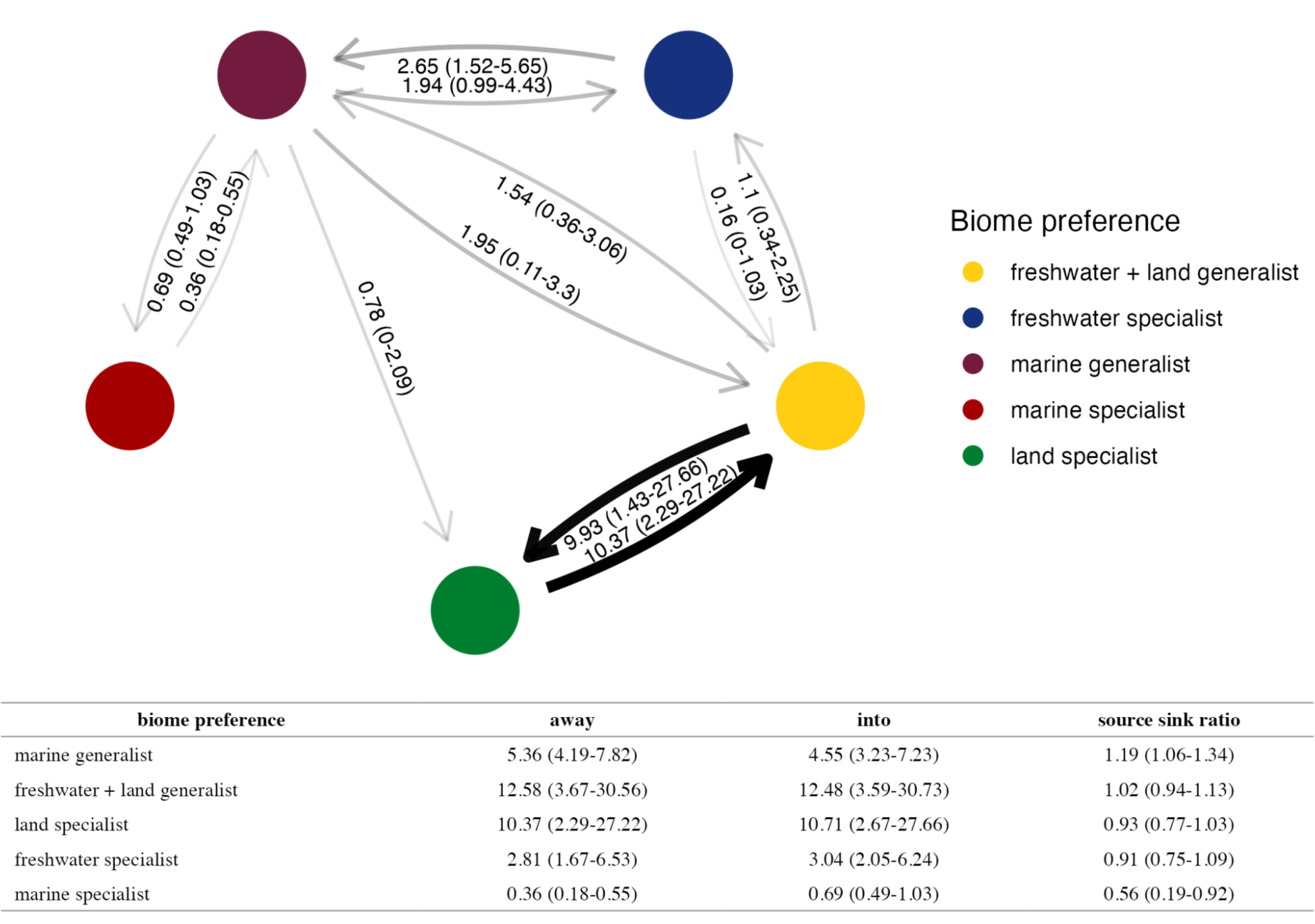
Bootstrapped (using stratified) transition rates between biome preferences for the best-supported model of discrete character evolution. The best-supported model was a simplification of the ARD where low transition rates were removed. The table shows the total rates into and away from each biome preference. Bootstrapping was done by randomly sampling tips within each biome preference such that each had the same number of tips in the tree. Mean estimates are presented, and values in brackets represent 95% confidence intervals. The size of the arrows is proportional to the transition rate. All values are labelled to two decimal places.

**Table S2.**
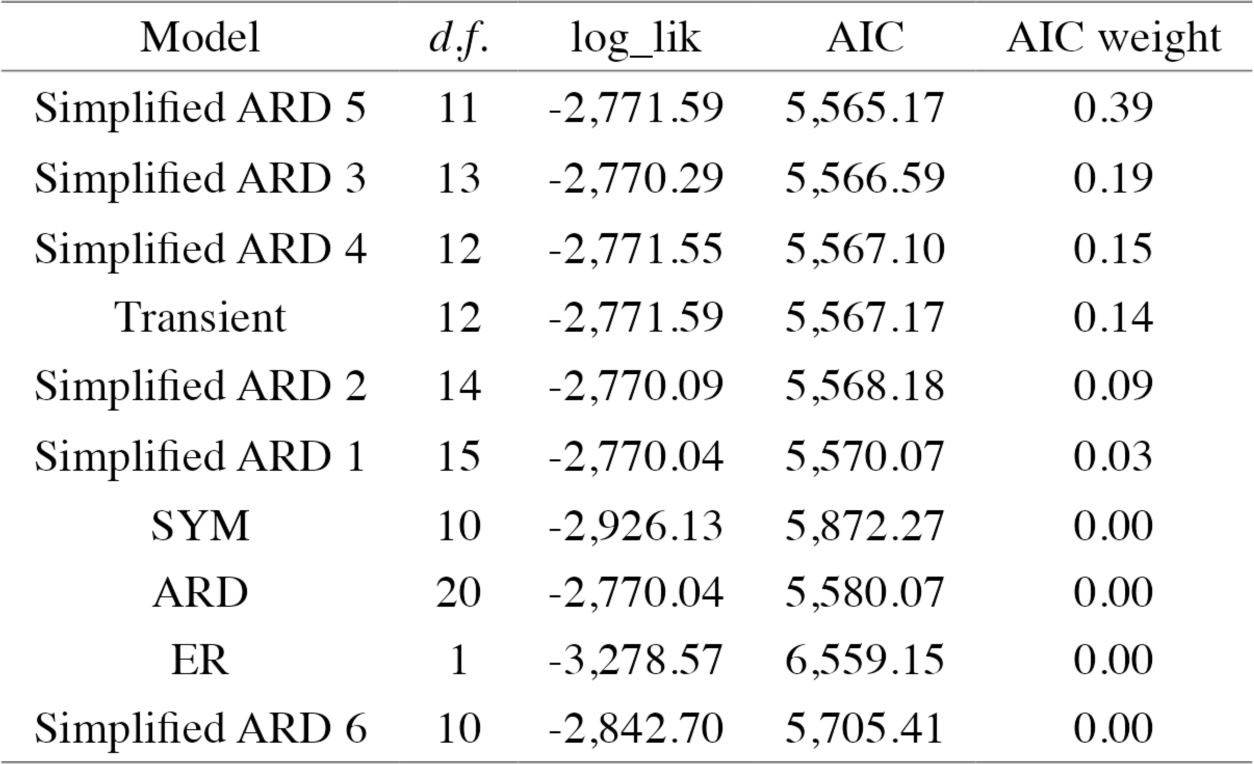
Model comparison of the Markov models exploring transition rates between biome preferences.

| Model | *d.f.* | log_lik | AIC | AIC weight |
| --- | --- | --- | --- | --- |
| Simplified ARD 5 | 11 | -2,771.59 | 5,565.17 | 0.39 |
| Simplified ARD 3 | 13 | -2,770.29 | 5,566.59 | 0.19 |
| Simplified ARD 4 | 12 | -2,771.55 | 5,567.10 | 0.15 |
| Transient | 12 | -2,771.59 | 5,567.17 | 0.14 |
| Simplified ARD 2 | 14 | -2,770.09 | 5,568.18 | 0.09 |
| Simplified ARD 1 | 15 | -2,770.04 | 5,570.07 | 0.03 |
| SYM | 10 | -2,926.13 | 5,872.27 | 0.00 |
| ARD | 20 | -2,770.04 | 5,580.07 | 0.00 |
| ER | 1 | -3,278.57 | 6,559.15 | 0.00 |
| Simplified ARD 6 | 10 | -2,842.70 | 5,705.41 | 0.00 |

**Table S3.** Model comparison of multi-state and concealed trait diversification rate models at different estimated sampling fractions.

| Sampled fraction | Model | Number of estimated parameters | Log Likelihood | AIC | AIC weight |
| --- | --- | --- | --- | --- | --- |
| 0.75 | CTD4 | 17 | -1,265.23 | 2,564.46 | 1.00 |
|  | CTD3 | 10 | -1,285.37 | 2,590.75 | 0.00 |
|  | MuHiSSE | 13 | -1,308.06 | 2,642.12 | 0.00 |
|  | CTD2 | 5 | -1,344.53 | 2,699.07 | 0.00 |
|  | MuSSE | 6 | -1,688.56 | 3,389.11 | 0.00 |
| 0.50 | CTD3 | 10 | -1,248.55 | 2,517.09 | 0.82 |
|  | CTD4 | 17 | -1,243.07 | 2,520.15 | 0.18 |
|  | MuHiSSE | 13 | -1,269.51 | 2,565.01 | 0.00 |
|  | CTD2 | 5 | -1,305.93 | 2,621.85 | 0.00 |
|  | MuSSE | 6 | -1,660.88 | 3,333.75 | 0.00 |

